# *LLS-SevEst - Late leaf spot severity estimator*. A machine learning approach to assessing *Nothopassalora personata* in peanut

**DOI:** 10.1101/2025.05.07.652206

**Authors:** T.A. Herrador, J. Migotti Scaglia, J.A. Paredes, L.I. Cazón

## Abstract

Late leaf spot (LLS), caused by *Nothopassalora personata*, is the most damaging foliar disease in peanut production worldwide, leading to significant yield losses if not properly managed. Accurate disease severity assessment is crucial for evaluating fungicide efficacy and implementing effective management strategies. This study aimed to develop and validate an automated image analysis model, *LLS-SevEst*, for quantifying LLS severity in peanut leaves. A dataset of 190 scanned leaf images was analyzed using three approaches: a fixed threshold-based segmentation, morphological preprocessing, and K-means clustering. Exploratory analyses revealed distinct brightness patterns between healthy and diseased tissues, guiding the development of classification functions. The threshold-based model yielded high false positive rates due to its inability to account for natural leaf variation, while the morphological preprocessing method improved segmentation marginally but still required manual adjustments. The K-means clustering approach achieved superior segmentation by objectively differentiating healthy tissue, lesions, and background, and showed high potential for automated, reproducible disease severity estimation. Future work should focus on integrating deep learning and expanding the dataset to improve model robustness and adaptability to other foliar pathosystems.

## INTRODUCTION

Peanut production plays a crucial role in global agriculture, serving as both an economic driver and a key raw material for the food industry. In Argentina, peanut cultivation is among the most important regional economies, with over 70% of production concentrated in Córdoba (Barberis 2023; Cámara Argentina del Maní 2022). During the 2023/24 season, approximately 300,000 hectares were cultivated, generating over $1 billion in foreign exchange earnings. However, peanut production faces significant phytosanitary challenges, with late leaf spot (LLS), caused by *Nothopassalora personata* (Berk. & M.A. Curtis), being the most damaging disease worldwide (Giordano et al. 2021). Under favorable conditions (95% relative humidity, ∼18°C), and without effective field management, LLS can cause severe yield losses (Shokes & Culbreath, 1997). The disease manifests as dark leaf spots surrounded by yellow halos, which, in severe cases, coalesce and drastically reduce photosynthetic efficiency. Lesions can also affect pegs, leading to pod detachment during harvest, further exacerbating yield losses (Cazón et al. 2025, Marinelli & March 2005; Oddino et al., 2018).

Integrated disease management primarily relies on fungicide applications, with strobilurins, triazoles, carboxamides, and chlorothalonil being the most widely used. Initial sprays are typically applied around 75 days after sowing or upon symptom detection, with subsequent applications depending on cultivar susceptibility, environmental conditions, and crop cycle duration (Giordano et al., 2021; Monguillot et al., 2023; Pedelini, 2021). Evaluating fungicide efficacy requires accurate disease severity assessment, which remains predominantly visual despite the availability of advanced techniques. While visual assessment is cost-effective, it is inherently subjective and influenced by evaluator expertise and fatigue. Additionally, pathosystem-related factors such as lesion size, location, and coalescence contribute to estimation errors that may lead to misinformed management decisions (Bock et al. 2020; Cazón et al. 2025; Del Ponte et al. 2021).

Various tools have been developed to improve visual assessments, including online training systems and diagrammatic scales. Recently, the first validated LLS standard area diagram was introduced (Cazón et al. 2025), and peanut leaf images were incorporated into Trainer2, an online training system (Del Ponte, 2023). Smart agriculture technologies, particularly multispectral and hyperspectral imaging, offer promising alternatives for disease severity quantification assessment (Chen et al., 2019; Omran, 2016). However, adoption remains limited due to operational complexity and costs. In contrast, RGB imaging with deep learning has shown high accuracy in peanut foliar disease detection, yet no validated tool currently exists for LLS severity quantification (Xu et al. 2023).

Given this context, this study aimed to develop and validate an automated image analysis tool in Python to significantly improve the accuracy and objectivity of peanut late leaf spot severity quantification. Additionally, the proposed model was compared with existing assessment methods.

## MATERIAL AND METHODS

### Image acquisition

To reach the objective, a total of 190 leaves with varying disease severity were collected in April 2023 from plants grown under controlled conditions at IPAVE-CIAP, Córdoba (Latitude: −31.46895, Longitude: −64.14730). The abaxial leaf surface was scanned using a CanoScan LIDE 300 flatbed scanner at 300 dpi. Shadows were removed using *Adobe Photoshop software* (2011), and 50 representative images were selected for model development. Additionally, the *pliman* package (Olivoto, 2022) in *R* (R Core Team, 2022) was used for comparison, applying segmentation techniques to distinguish healthy from diseased tissue.

### Image processing

Image processing was conducted using *Python 3*.*x* (Python Software Foundation, 2023) within *Jupyter Notebooks* (Kluyver et al., 2016) on Google Colab (https://colab.research.google.com). Exploratory analyses were conducted to identify significant differences in pixel luminosity between healthy and diseased areas. Several image filters were applied to enhance contrast, and histogram plots were generated to visualize brightness intensity distributions. For this, we used Pandas (McKinney, 2023), Numpy (Numpy, 2023), OpenCV (Itseez, 2023), and Matplotlib (Hunter, 2023). Once these patterns were identified, three approaches were employed to estimate the percentage of leaf area affected by *N. personata*:

1. Threshold-Based Model: A classification function was developed to distinguish pixels belonging to healthy and diseased areas using a fixed intensity threshold of 80. This value was determined by analyzing the pixel brightness distribution through image histograms. Pixels with intensity values below 80 were classified as lesions, while those above were considered healthy tissue. Disease severity was calculated as the proportion of lesion pixels relative to the total leaf area.
2. Morphological Preprocessing Model: A morphological operation consisting of erosion followed by dilation (using a small 3×3 elliptical structuring element) was applied to improve segmentation. This modification reduced misclassifications caused by natural variations in leaf coloration. Additionally, an interactive feature allowed users to manually adjust the threshold based on image histograms, enhancing adaptability and accuracy.
3. K-Means Clustering Model: The *scikit-learn* library (Pedregosa et al., 2011) was used to implement K-means clustering with three predefined clusters: background, healthy tissue, and lesions. Images were converted into RGB matrices, smoothed to reduce noise, and transformed into a dataset where each pixel was represented by its position and color channel values. Additional features were generated from color differences (e.g., red-blue, red-green, blue-green), and data were normalized using StandardScaler (Pedregosa et al., 2011). The K-means model was executed with three clusters and ten iterations. Disease severity was then calculated as the proportion of pixels classified as lesions relative to total leaf area.

## RESULTS AND DISCUSSION

### Exploratory Analysis

Grayscale conversion effectively distinguished healthy and diseased areas based on pixel luminosity (Fig. 1A). Histogram analysis revealed a distinct intensity peak corresponding to the background (255 intensity units). In contrast, leaf areas (healthy + diseased) displayed a Gaussian distribution (Fig. 1B).

**Figure 1.**
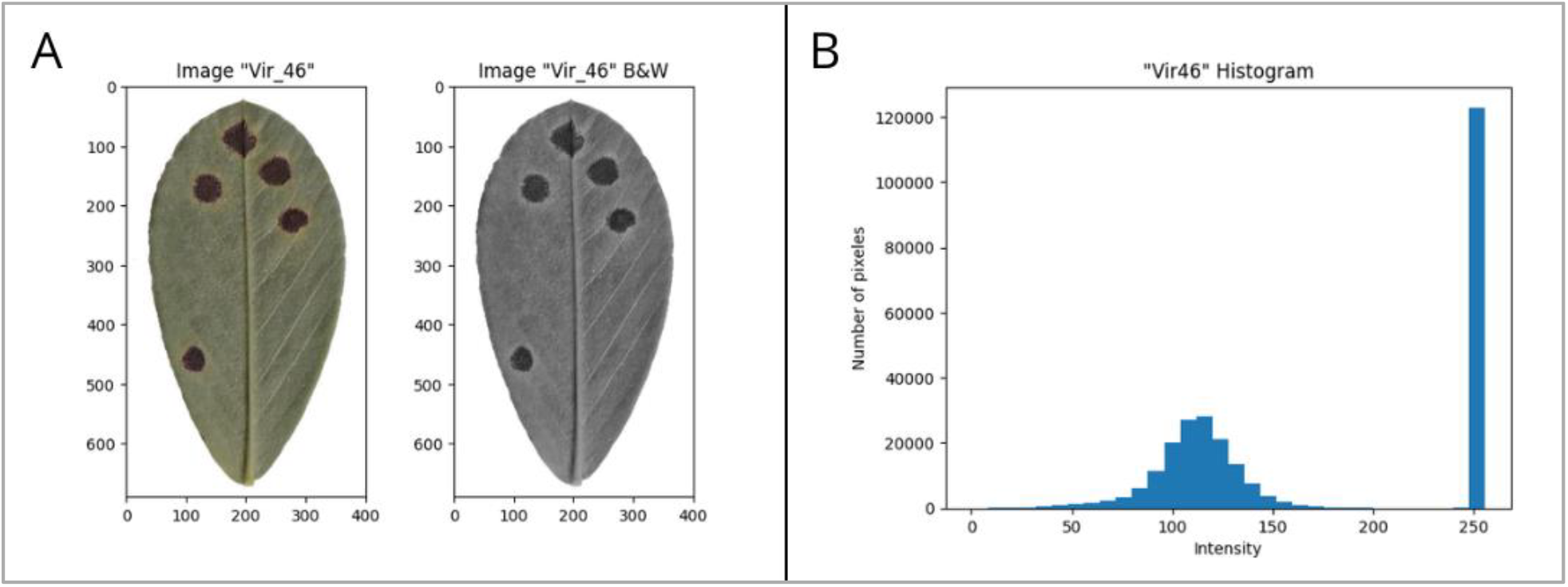
**A**: Comparison between the original image and the grayscale image, converted using the color_rgb2gray function from the *cv2* library. **B**: Histogram of the pixel luminance in the grayscale image.

Further analysis showed different intensity patterns between healthy and diseased regions (Fig. 2). Healthy tissue exhibited a Gaussian distribution (Fig. 2B), while lesions displayed bimodal distributions due to overlapping brightness intensities at lesion margins (Fig. 2A, C).

**Figure 2.**
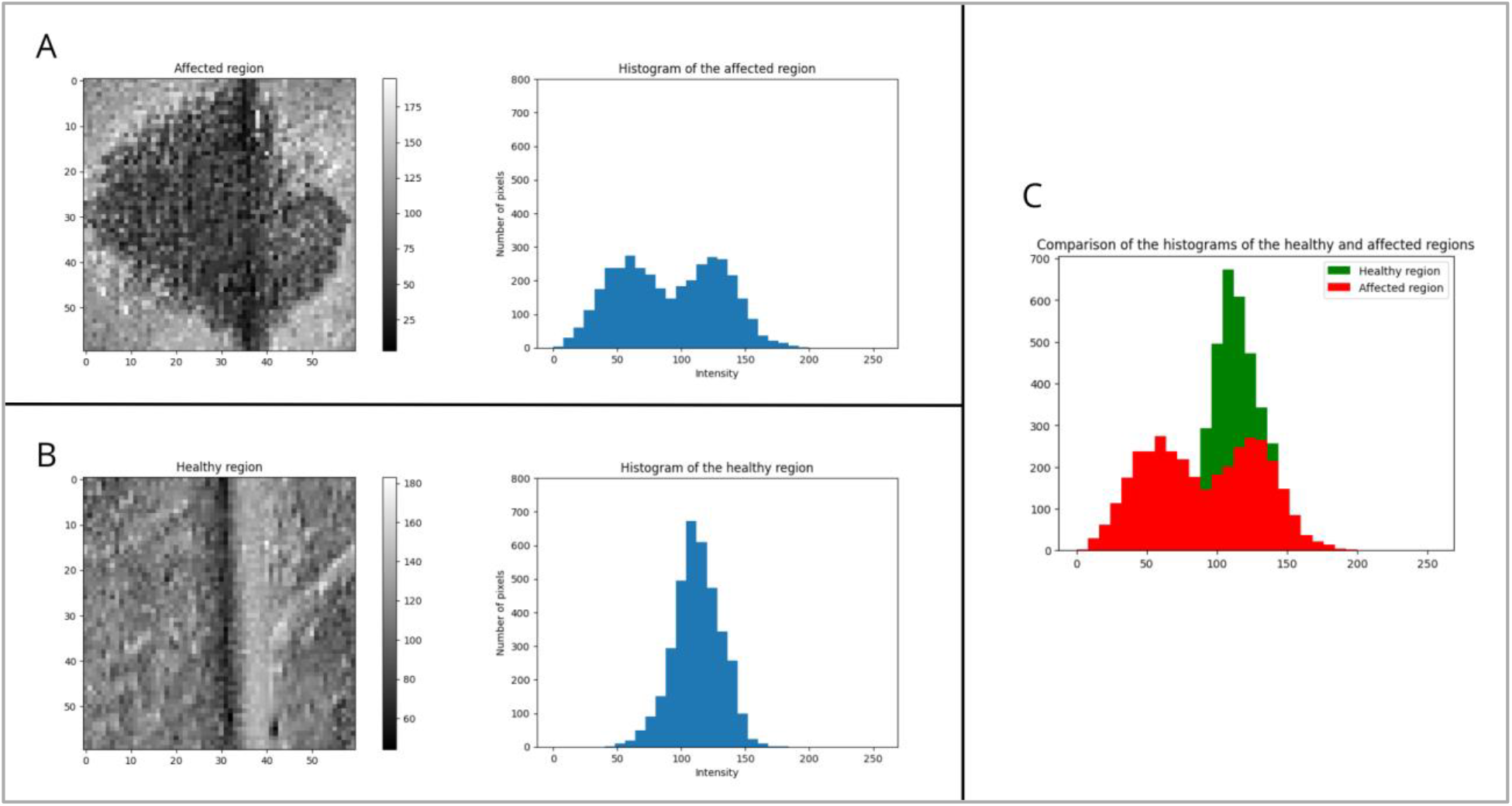
**A**: Cropped grayscale image of the lesion area caused by *Northopasalora personata*, alongside the histogram of pixel luminance in this region. **B**: Cropped grayscale image of a non-lesioned area, alongside the histogram of pixel luminance in this region. **C**: Superimposition of the generated histograms, with pixels from the lesion area represented in red and pixels from the healthy area represented in green.

### Model performance evaluation

For the Threshold-based model, a threshold of 80 grayscale intensity units (ranging from 0 for black to 255 for white) was set for the classification of different areas. Pixels below this threshold were classified as lesions, while those above were considered healthy leaf tissue. However, this approach failed to accurately differentiate between healthy and diseased areas, resulting in a high rate of false positives. One example of this misclassification was the identification of shadows cast by the leaf’s midrib as diseased areas (Fig. 3A). This limitation led to considerable variability when comparing the severities obtained with this model and those calculated using pliman (Fig. 3B). Although this approach is conceptually valid (Barbedo 2013, 2016), the results suggest that segmentation based on a fixed luminosity threshold is insufficient for accurately distinguishing between healthy and lesioned areas, particularly in leaves with natural variations in brightness and color in this pathosystem. The function was later modified to allow manual threshold adjustment, enabling better adaptation to the specific characteristics of each leaf image.

**Figure 3.**
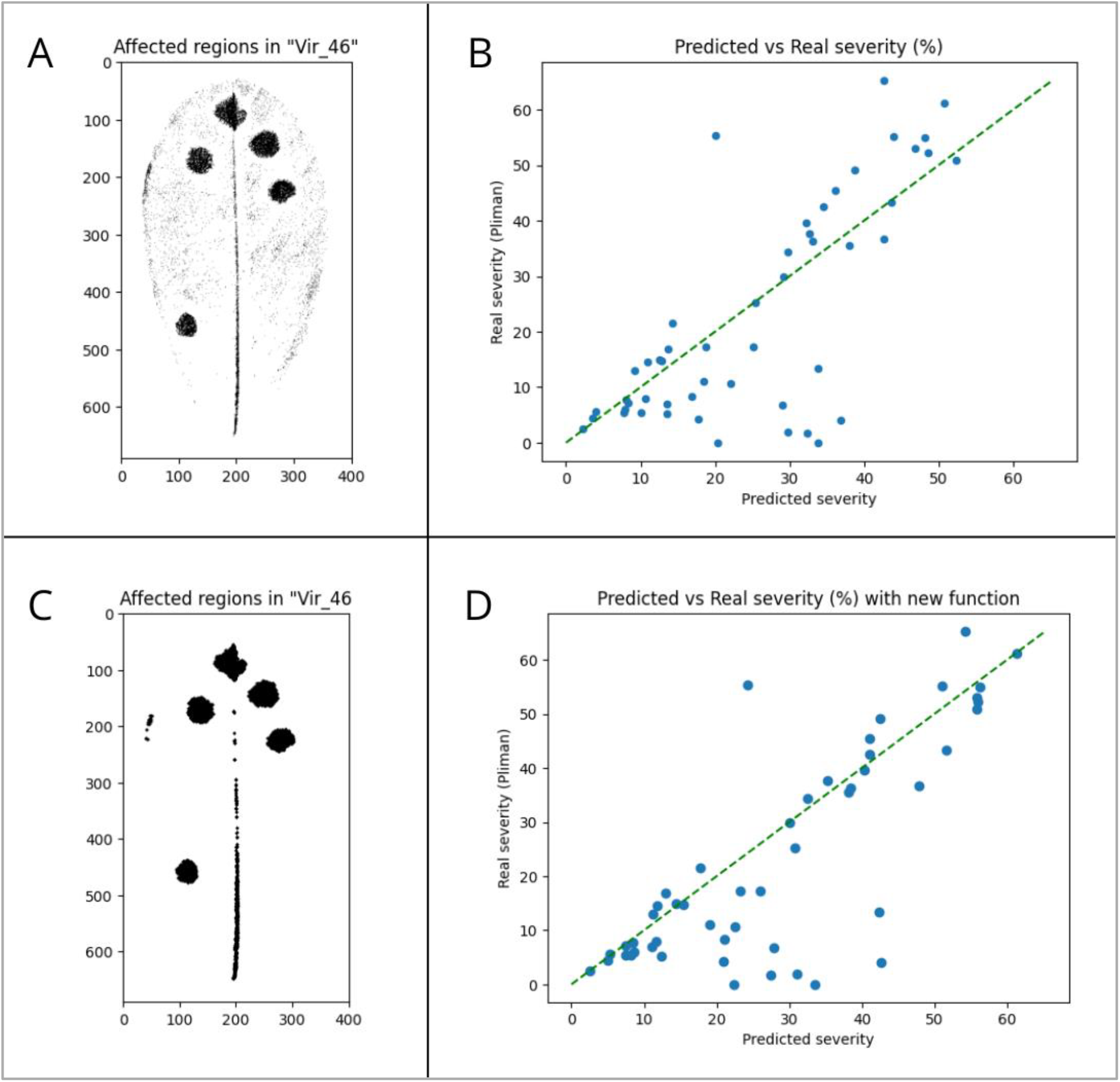
**A**: Segmentation performed by the first generated model, based on thresholding pixel luminance. **B**: Comparative scatter plot between the severity percentage predicted by the model and the one previously calculated using *pliman*. **C**: Segmentation performed by the modified model, incorporating morphological operations of erosion and dilation. **D**: Comparative scatter plot between the severity percentage predicted by the modified model and the one previously calculated using pliman.

Regarding the Morphological Preprocessing Model, the dilation followed by erosion function helped “smooth” the images, reducing some false positives (Gonzalez & Woods, 2018). However, darker-toned areas remained undistinguished (Fig. 3C), still causing significant dispersion in the severity estimates when compared to *pliman* (Fig. 3D). These results suggest that a fixed threshold cannot be universally applied and that manual adjustments are required for each specific case, which reduces the practicality and automation of the method. It is important to note that achieving accurate segmentation remains a significant challenge in image-based automatic plant disease identification (Barbedo, 2016).

Applying the K-means model enabled effective image segmentation, accurately distinguishing healthy tissue, affected areas, and background regions. This approach was used by Phadikar et al. (2012) for rice disease classification. The cluster visualization revealed a clear distinction between *N. personata*-affected areas and healthy tissues, allowing for an objective assessment of disease severity (Fig. 4A). The model was applied to all images in the dataset, enabling the calculation of the affected area percentage in each case.

**Figure 4.**
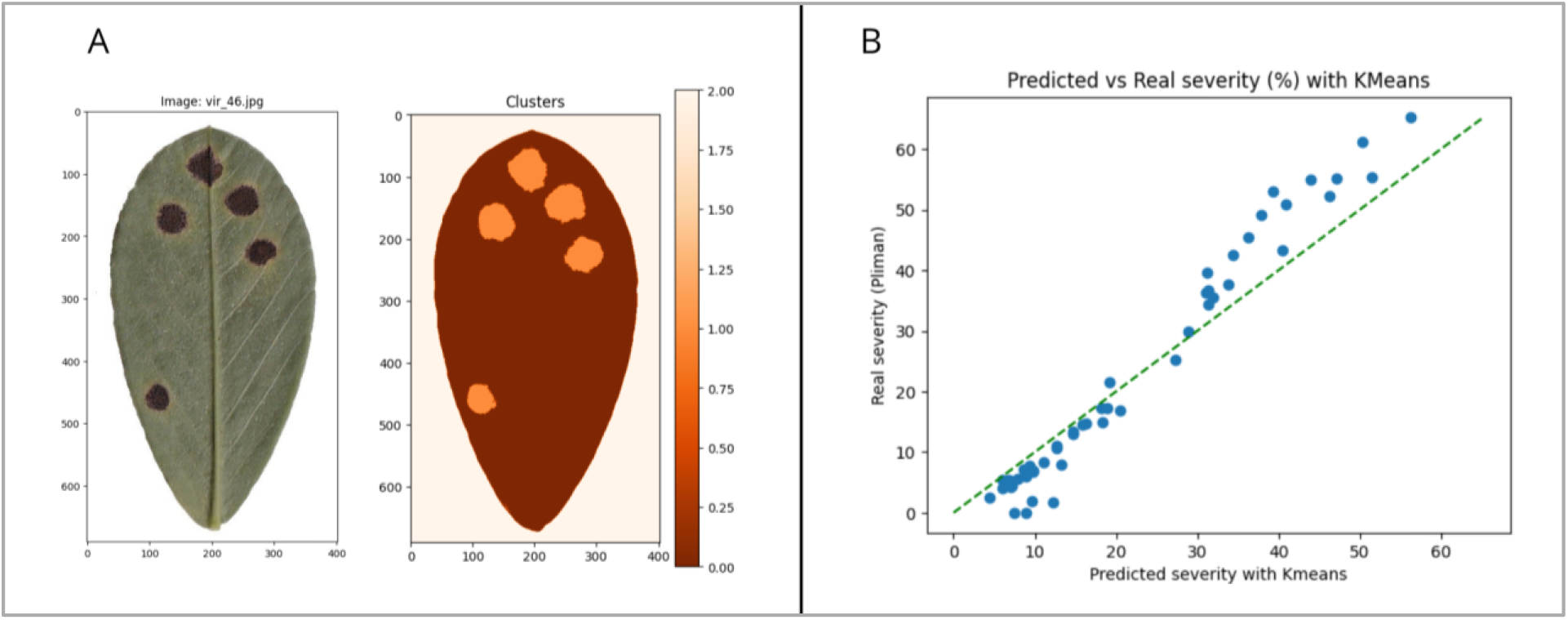
**A**: Comparison between the original image and the image segmented by the K-Means clustering model, where white corresponds to the “background” cluster, orange to the “healthy” cluster, and red to the “lesion” cluster. **B**: Comparative scatterplot showing the severity percentage results obtained by the K-Means model and those previously calculated using *pliman*.

At first glance, a significant improvement is observed compared to the results obtained with the initial functions. However, discrepancies remain between the previously calculated severity and the severity estimated using the developed model (Fig. 4B). When images with considerable discrepancies between *pliman* and the *LLS-SevEst* model were closely analyzed, it was observed that some photosynthetic regions were not classified as lesions by pliman (Fig. 5). Since many authors emphasize the high efficiency of pliman in determining plant severity, a methodological error likely exists when creating the palettes corresponding to the initial image processing stages using *pliman*. This can lead to errors in the overall classifications (Del Ponte 2023).

**Figure 5.**
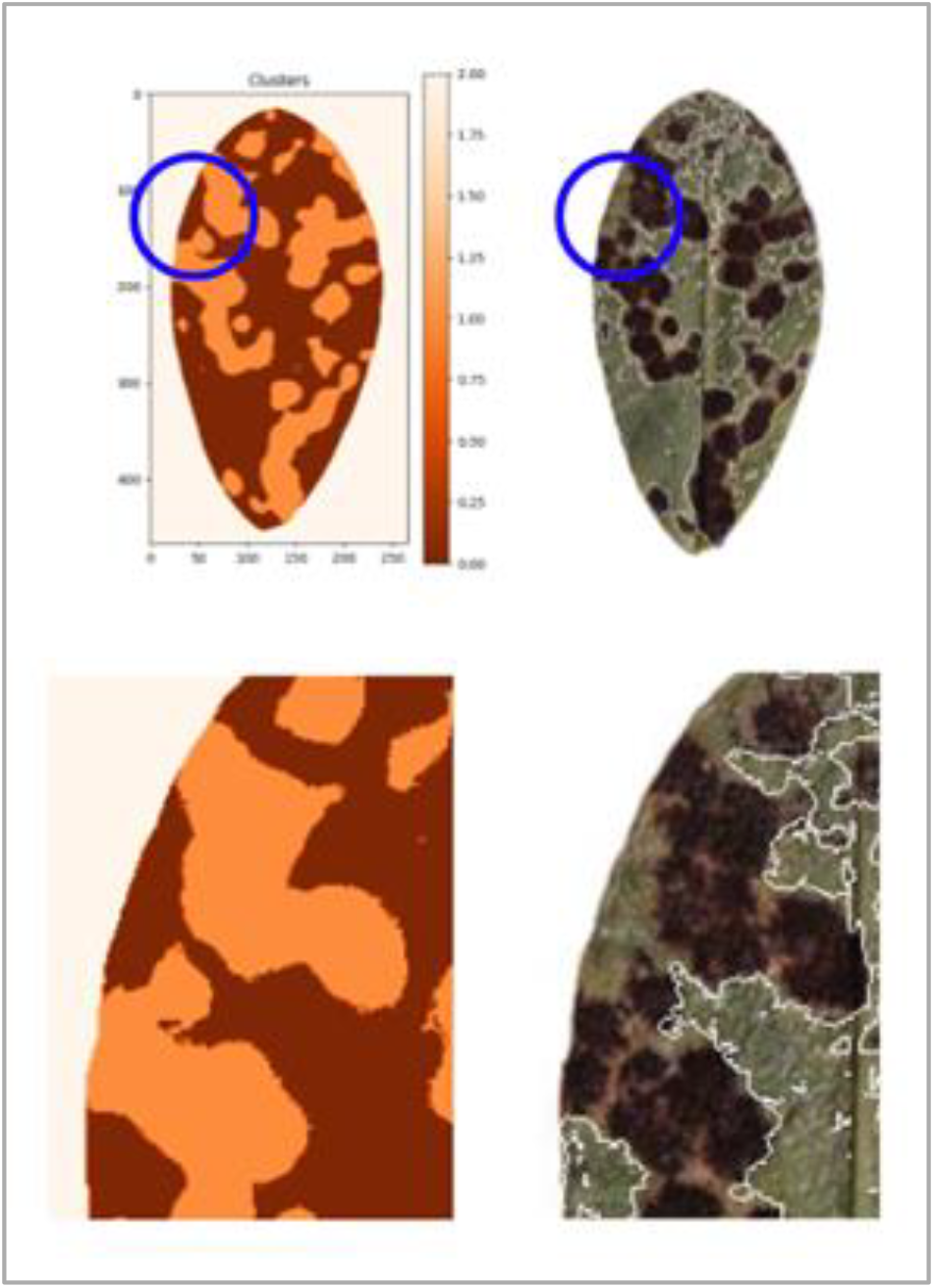
Comparison between the segmentation generated using K-Means clustering and the segmentation obtained with *pliman*. The region marked with a blue circle is magnified to visualize better the discrepancies between the segmentations produced by both models.

**Figure 6.**
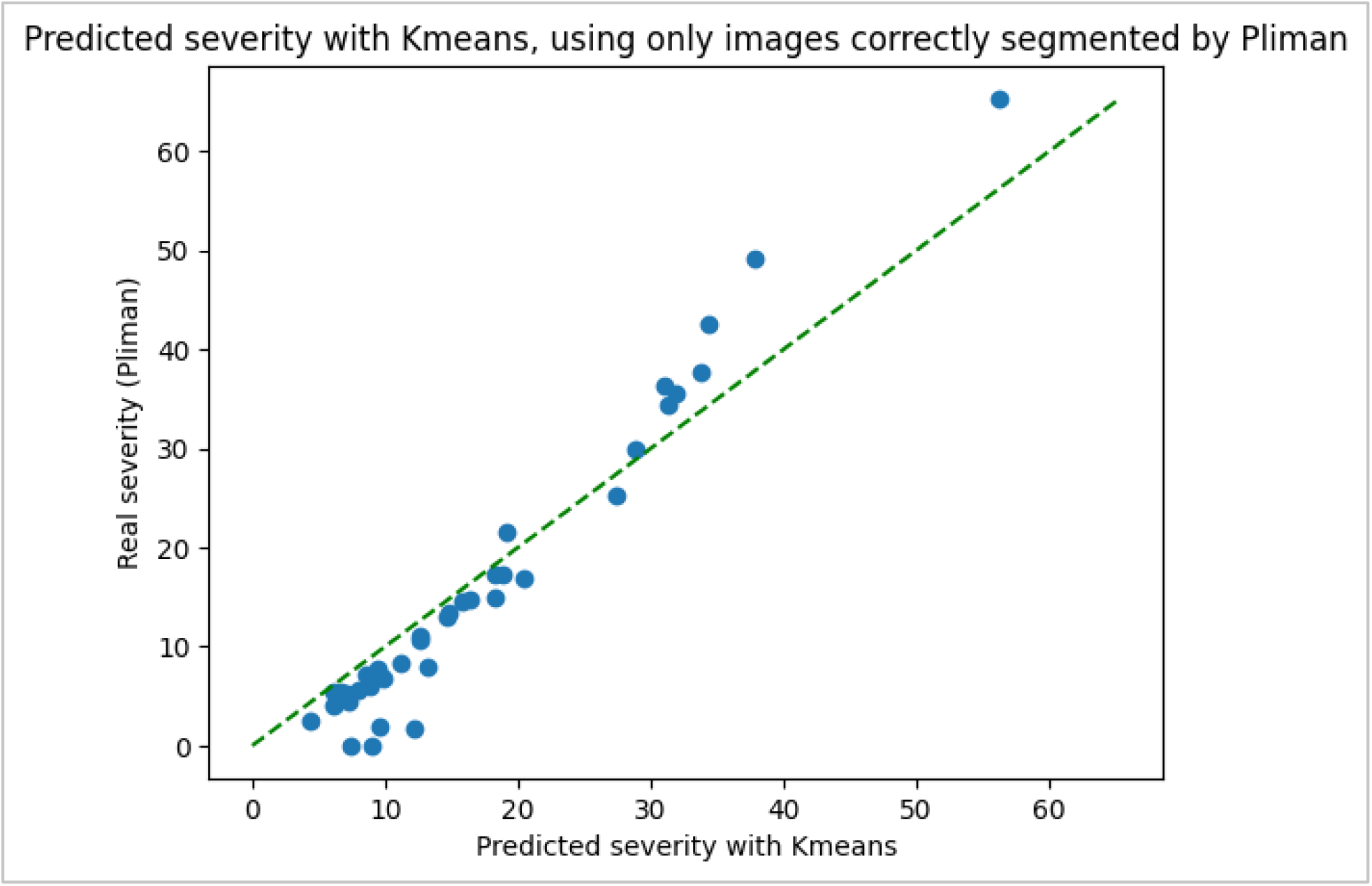
Comparative scatter plot between the severity percentage predicted by KMeans and the one previously calculated using *pliman*. This time, only with images correctly segmented by Pliman.

When images exhibiting this discrepancy were excluded, the model fit much better, suggesting that K-means is an efficient tool for image segmentation in quantifying foliar diseases, facilitating the automatic evaluation of peanut leaf spot severity.

Future research should focus on improving the model by integrating deep learning techniques, such as neural networks, and incorporating larger and more diverse datasets (Ferentinos 2018; Mohanty et al. 2016). Additionally, flexibility in threshold selection and the ability to perform manual adjustments remain crucial factors for ensuring accuracy and practical applicability across different contexts and leaf types. Lastly, the *LLS-SevEst* model shows promising results, demonstrating the potential of machine learning approaches for plant disease severity assessment. With further refinements and optimizations, it could be applied not only to peanut late leaf spot but also to other plant diseases, expanding its usefulness and relevance in plant pathology.

## ACKNOWLEDGEMENTS

Luis Ignacio Cazón and Juan Andrés Paredes want to thank INTA and the Fundación Maní Argentino for the support. Tiago Alejo Herrador and Juliana Migotti Scaglia are thankful to Rodrigo Parola for facilitating the initial contact between them and INTA, which was key to establishing the connection that made this research possible.

## DATA AVAILABILITY

The datasets generated during and/or analyzed during the current study are available from the corresponding author on reasonable request.

## CONFLICTS OF INTEREST

All authors declare that they have no conflicts of interest.

## Notes

### Competing Interest Statement

The authors have declared no competing interest.

## REFERENCES

Adobe Photoshop. 2011. (versión 12.1) Adobe Inc. https://www.adobe.com/products/photoshop.html

Barbedo, J. G. A. 2013. Digital image processing techniques for detecting, quantifying, and classifying plant diseases. SpringerPlus, 2013, Vol. 2, No. 660

Barbedo, J. G. A. 2016. A review on the main challenges in automatic plant disease identification based on visible range images. Biosystems Engineering, 144, 52–60. 10.1016/j.biosystemseng.2016.01.017xz.

Barberis, N. A. 2023. Análisis de la evolución de indicadores económicos para el cultivo de maní, provincia de Córdoba, campañas 2010/11-2022/23. In 38º Jornada Nacional de Maní.

Bock, C. H.; Barbedo, J. G. A.; Del Ponte, E. M.; Bohnenkamp, D.; Mahlein, A.-K. 2020. From visual estimates to fully automated sensor-based measurements of plant disease severity: Status and challenges for improving accuracy. Phytopathology Research, 2, 9.

Cámara Argentina del maní. 2022. Clúster manisero. Available online: https://camaradelmani.org.ar/cluster-manisero. verified: April 25th 2025.

Cazón, L.I.; Paredes, J.A.; González, N.R.; Conforto, E. C.; Suarez, L.; Del Ponte, E. M. 2025. Optimizing visual estimation of peanut late leaf spot severity with online training sessions and standard area diagrams. Eur J Plant Pathol 10.1007/s10658-025-03016-1

Chen, T.; Zhang, J.; Chen, Y.; Wan, S.; Zhang, L. 2019. Detection of peanut leaf spots disease using canopy hyperspectral reflectance. Computers and Electronics in Agriculture, 156, 677–683.

Del Ponte, E. M.; Cazón, L. I.; Alves, K., Pethybridge, S.; Bock, C. 2021. How much do standard area diagrams improve accuracy of visual estimates of plant disease severity? A systematic review and meta-analysis. Tropical Plant Pathology. Avance online publication. 10.1007/s40858-021-00479-5.

Del Ponte, E. M. 2023. Training sessions. In R for plant disease epidemiology (R4PDE). Author. https://r4pde.net. xVerified: March 26th 2025.

Ferentinos, K. P. 2018. Deep learning models for plant disease detection and diagnosis. Computers and Electronics in Agriculture, 145, 311–318.

Giordano, D. F.; Pastor, N.; Palacios, S.; Oddino, C. M.; Torres, A. M. 2021. Peanut leaf spot caused by Nothopassalora personata. Tropical Plant Pathology, 46, 139151.

Gonzalez, R. C. and Woods, R. E. 2018. Digital Image Processing (4th ed.). Pearson.

Hunter, J. D. 2023. Matplotlib [Software]. Retrieved from <https://matplotlib.org/>

Itseez. 2023. OpenCV [Software]. Retrieved from <https://opencv.org/>

Kluyver, T.; Ragan-Kelley, B.; Pérez, F.; Granger, B. E.; Bussonnier, M.; Frederic, J.; Willing, C. 2016. Jupyter Notebooks – a publishing format for reproducible computational workflows. In F. Loizides & B. Schmidt (Eds.), Positioning and Power in Academic Publishing: Players, Agents and Agendas (pp. 87–90). IOS Press. 10.3233/978-1-61499-649-1-87

Marinelli, A. and March, G. J. 2005. Viruela. In A. Marinelli & G. J. March (Eds.), Enfermedades del maní en Argentina (pp. 13–39). Ediciones Biglia.

McKinney, W. 2023. Pandas [Software]. Retrieved from <https://pandas.pydata.org/>

Mohanty, S. P.; Hughes, D. P.; Salathé, M. 2016. Using deep learning for image-based plant disease detection. Front. Plant Sci. 7:1419. doi: 10.3389/fpls.2016.01419

Monguillot, J. H.; Bernardi Lima, N.; Paredes, J. A.; Giordano, D. F.; Oddino, C.; Rago, A. M.; Carmona, M.; Conforto, E. C. 2023. Caracterización de aislados de Nothopassalora personata agente causal de la viruela tardía del maní. In 38º Jornada Nacional de Maní.

NumPy Development Team. 2023. NumPy [Computer software]. https://numpy.org

Oddino, C.; Giordano, F.; Paredes, J.; Cazón, L.; Giuggia, J.; Rago, A. 2018. Efecto de nuevos fungicidas en el control de viruela del maní y el rendimiento del cultivo. Ab Intus, 1, 9–17.

Olivoto, T. 2022. Lights, camera, pliman! An R package for plant image analysis. Methods in Ecology and Evolution, 13(4), 789–798.

Omra, E. S. E. 2016. Early sensing of peanut leaf spot using spectroscopy and thermal imaging. Archives of Agronomy and Soil Science, 63, 883–896.

Pedelini, R. 2021. MANÍ: Guía práctica para su cultivo. FMA press.

Pedregosa, F.; Varoquaux, G.; Gramfort, A.; et al. 2011. Scikit-learn: Machine Learning in Python. Journal of Machine Learning Research, 12, 2825–2830.

Phadikar, S.; Sil, J.; Nayak, J. 2012. Rice diseases classification using feature selection and rule generation techniques. Computers and Electronics in Agriculture, 90, 76–85.

Python Software Foundation. 2023. Python Language Reference (Version 3.x). https://www.python.org

R Core Team. 2022. R: A Language and Environment for Statistical Computing. R Foundation for Statistical Computing, Vienna, Austria. https://www.R-project.org/. verified: February 10th 2024.

Shokes, F. M. and Culbreath, A. K. 1997. Early and late leaf spots. In: Kokalis-Burelle N, Porter, D.M.; Rodríguez-Kábana, R.; Smith D.H.; Subrahmanyam, P. (Eds) Compendium of peanut diseases, 2nd ed. APS Press, pp 17–20.

Xu, L.; Cao, B.; Ning, S.; et al. 2023. Peanut leaf disease identification with deep learning algorithms. Mol Breeding 43, 25. 10.1007/s11032-023-01370-8

